# Versatile Product Detection via Coupled Assays for Ultra-high-throughput Screening of Carbohydrate-Active-Enzymes in Microfluidic Droplets

**DOI:** 10.1101/2023.03.29.534725

**Authors:** Simon Ladeveze, Paul J. Zurek, Tomasz S. Kaminski, Stephane Emond, Florian Hollfelder

## Abstract

Enzyme discovery and directed evolution are the two major contemporary approaches for the improvement of industrial processes by biocatalysis in various fields. Customization of catalysts for improvement of single enzyme reactions or de novo reaction development is often complex and tedious. The success of screening campaigns relies on the fraction of sequence space that can be sampled, whether for evolving a particular enzyme or screening metagenomes. Ultrahigh-throughput screening (uHTS) based on in-vitro compartmentalization in water-in-oil emulsion of picolitre droplets generated in microfluidic systems allows screening rates >1 kHz (or >107 per day). Screening for Carbohydrate Active Enzymes (CAZymes) catalysing biotechnologically valuable reactions in this format presents an additional challenge, because the released carbohydrates are difficult to monitor in high throughput. Activated substrates with large optically active hydrophobic leaving groups provide a generic optical readout, but the molecular recognition properties of sugars will be altered by incorporation of such fluoro- or chromophores and their typically higher reactivity, as leaving groups with lowered pKa values compared to native substrates make observation of promiscuous reactions more likely. To overcome these issues, we designed microdroplet assays in which optically inactive carbohydrate products are made visible by specific cascades: the primary reaction of an unlabelled substrate leads to an optical signal downstream. Successfully implementing such assays at the picoliter droplet scale allowed us to detect glucose, xylose, glucuronic acid and arabinose as final products of complex oligosaccharide degradation by glycoside hydrolases by absorbance measurements. Enabling the use of uHTS for screening CAZyme reactions that have been thus far elusive will chart a route towards faster and easier development of specific and efficient biocatalysts for biovalorisation, directing enzyme discovery towards catalysts for their natural rather than model substrates.

**Graphical abstract / TOC:** 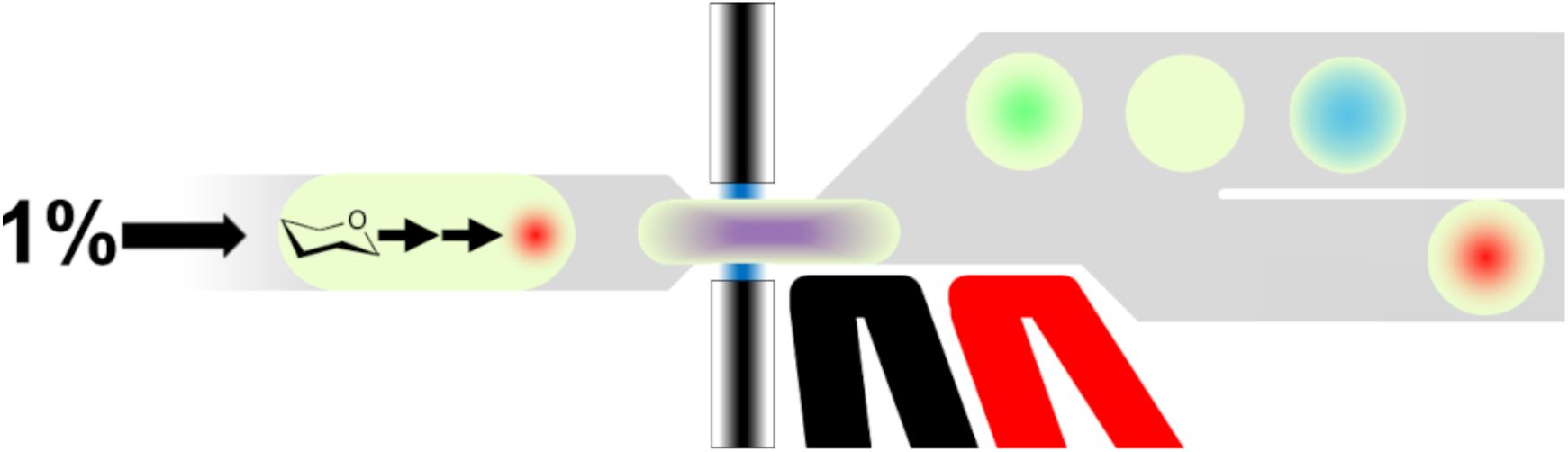

## INTRODUCTION

Carbohydrate-Active Enzymes (CAZymes) are a central class of enzymes with relevance for white biotechnology, *i*.*e*. sustainable chemistry for the food and feed industries, human health, material sciences or biofuels. A very large number of CAZymes are already in use but many more enzymes with defined specificities are needed in the post-fossil economy^1^. Biocatalysis plays an increasingly important role in the sustainable production of a variety of commodity products or for the generation of decarbonated biofuels. These eco-friendly industries rely on the use of renewable biomass feedstocks (such as cellulose and hemicelluloses, starch, chitin, etc) and their enzymatic deconstruction to simple monosaccharides prior to the conversion into products of interest.

In order to achieve these goals, new enzymatic catalysts must be found, either by discovery of unreported catalytic activities or by improvement or repurposing of currently used enzymes through directed evolution, often starting from empirically identified promiscuous activities. In both approaches screening of large numbers of enzyme variants in sequence space is essential and its success is determined by the experimental strategy employed.

In this regard, one of the most advanced technologies that combines both the highest throughput and the least reactant volume requirements is in-vitro compartmentalization (IVC). It enables the rapid screening of vast (>10^7^) libraries^2^ in microfluidic devices in monodisperse emulsion droplets with picoliter volumes. A wide range of reactions has been shown to be monitored in droplets^3^ (*e*.*g* hydrolases, esterases, proteases or oxidases)^4^, using both absorbance^5^ or fluorescence detection^6^. Discovery of CAZymes in droplets has been demonstrated in two directed evolution campaigns^7,8^, in one metagenomic screening^9,10^ and efforts towards further high-throughput screening systems have been made^11–13^. Many screening campaigns rely on fluorogenic or chromogenic model^10,14^, with hydrophobic leaving groups (*p*-nitrophenyl, 4-methylumbelliferyl)^12^ or solid colorimetric substrates (azo-dyed and azurine cross-linked polysaccharides) that are mostly used for the detection of specific activities on insoluble substrates, such as plant cell-wall polysaccharides^15^ and do not fully resemble natural sugar substrates. Other colorimetric methods for the detection of glycoside hydrolases (GHs) include the 3,5-dinitrosalicylic acid (DNS) assay^16^, which allows the detection of reducing sugars, or the Congo red method^17^ for detection of GHs degrading polysaccharides. Carbohydrate hydrolases and CAZymes in general are often assayed with such substrates for improvement of catalytic properties^18^, thermostability^19^ or for shifting their substrate specificity^20^. These substrates are also used for enzyme discovery in functional metagenomic screening campaigns^21^. However, these methods are not carbohydrate-specific and suffer from several limitations that prevent their use for CAZyme activity screening in droplets: (i) When considering the first rule of directed evolution ‘*you get what you select for’*, any selection method relying on modified substrates can lead to false positive recovery. Indeed, one cannot exclude that *p*NP- or 4-methylumbelliferyl-sugars recovered hits may be due to the affinity of the enzyme to the chromophore, directing the screening efforts in undesired reactions^22^. Also, the leaving group p*K*_a_ values of *p*NP- (~7.1) or 4-methylumbelliferyl (p*K*_a_ ~7.8) are orders of magnitude below that of a sugar leaving groups (p*K*_a_ ~12-14), rendering the reaction of model substrates thermodynamically much less demanding and thus easier to catalyze. (ii) Practically, droplet microfluidic campaigns can be compromised when hydrophobic product molecules leave the droplet they originated in, causing cross-contamination over time and making it hard to identify genuine hits over background. Indeed, para-nitrophenol or 4-methylumbelliferone have half-lives of less than one hour in droplets. Additional synthetic modification is possible, but can be difficult^23^. (iii) For each target sugar, fluorogenic or chromogenic substrates have to be synthesized. The attachment of a leaving group to a sugar is not difficult in principle, but given delocalization (*e*.*g*. in fluorescein) synthesis via nucleophilic attack can be inefficient. A one-fits-all system would remove worries about synthetic efficiency. (iv) Finally, colorimetric methods often used in plate assays require multiple incubation steps employing harsh chemicals and extreme pHs that are not compatible with cell growth in droplets, a step used to enhance sensitivity^24^.

Detection of reaction turnover in cascade reactions is an alternative: one or several accessory/coupling enzymes are employed to convert the primarily released carbohydrate generated by the enzyme of interest, leading to a monitorable by-product (*e*.*g*. NADH, Figure 1). Taking advantage of the modular structure of the microfluidic system, we describe a generic modular screening platform for the specific detection of various CAZyme activities (schematically shown in Figure 2). The versatility of this set-up is demonstrated by detecting the release of four types of monosaccharides (xylose, arabinose, glucose and glucuronic acid, Figure 3A-D) generated by degradation of plant cell-wall polysaccharides (cellulose and xylans) that can now be detected at ultrahigh throughput with a generic coupled assay in droplets.

**Figure 1:**
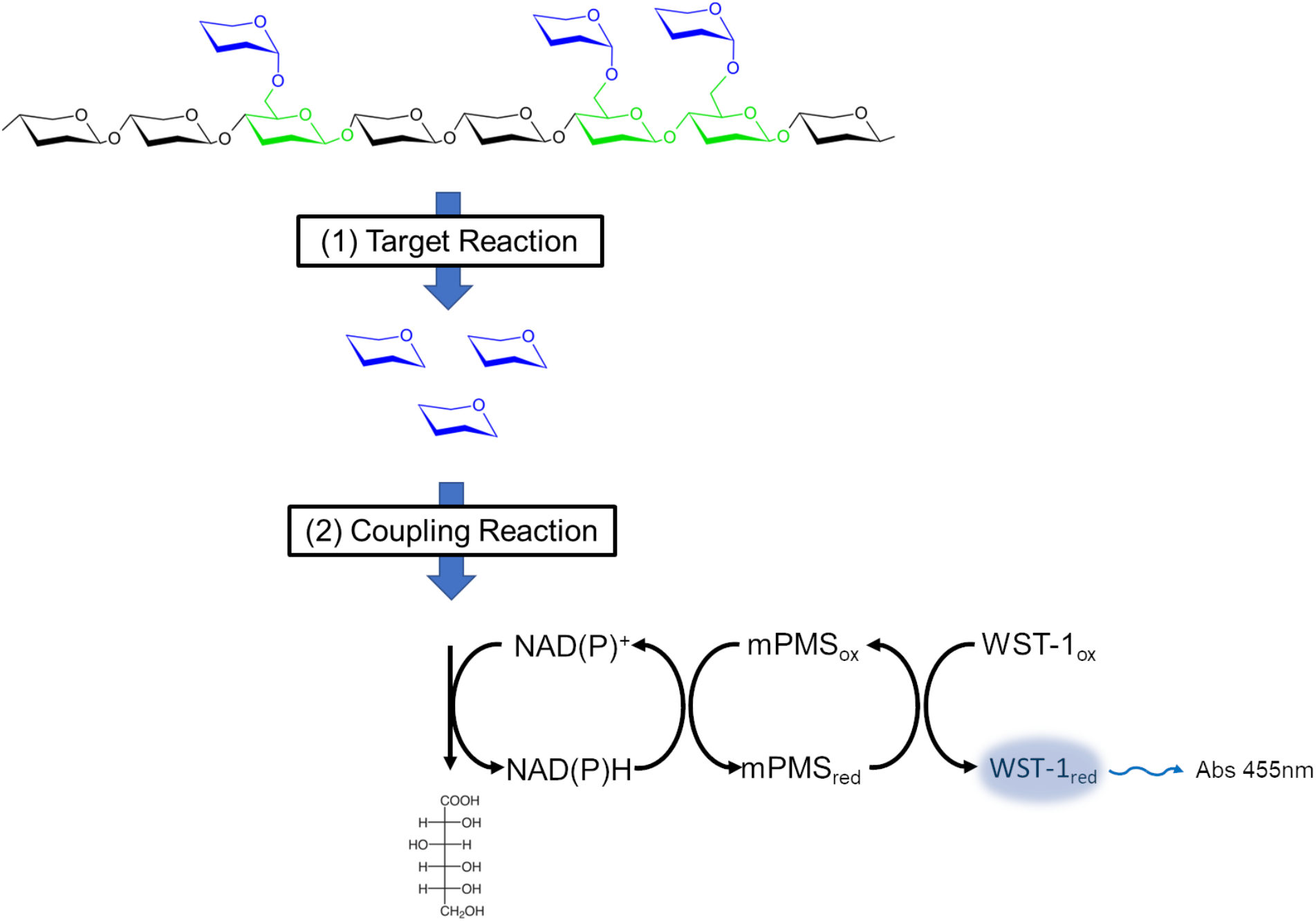
General principle of enzymatic cascades allowing the specific detection of carbohydrates. Different monosaccharide components of an oligosaccharide (depicted as coloured hexagons) make up a natural substrate, the degradation of which cannot directly monitored by optical means. When the targeted enzymatic activity releases (**1**) specific carbohydrates (blue hexagons) from a natural substrate their emergence is detected with a specific coupling reaction (or alternatively a cascade of coupling reactions) (**2**). The coupling system generates a product (blue halo) that can be readily monitored by its absorbance or fluorescence.

**Figure 2:**
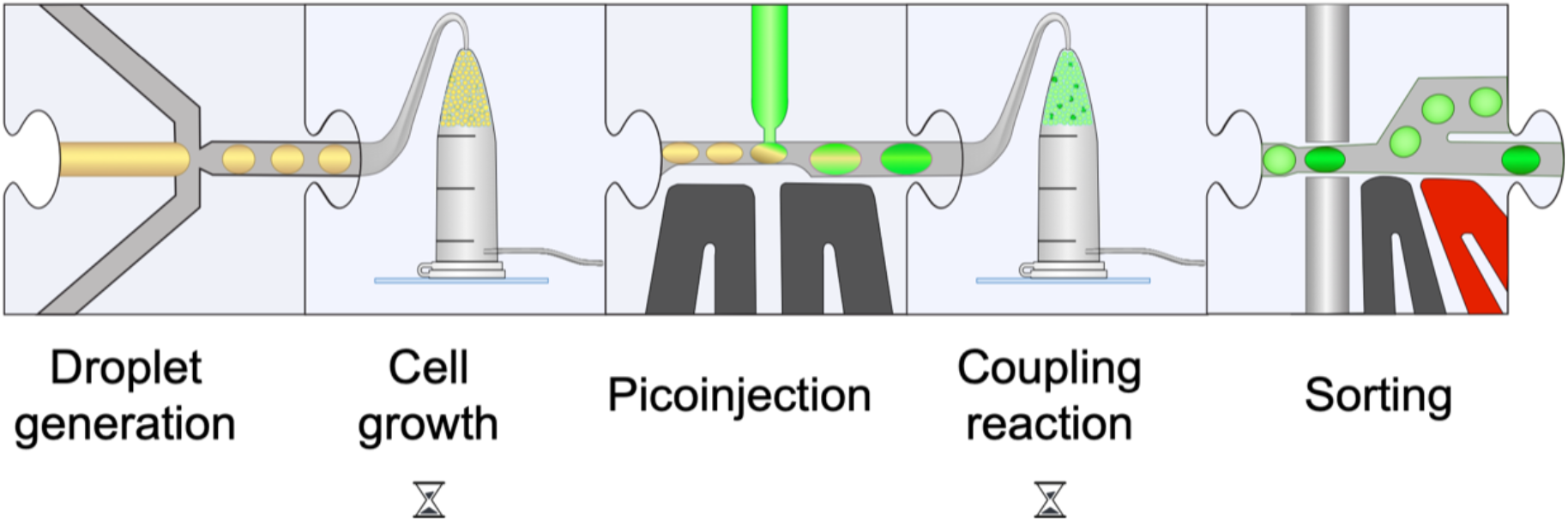
Modular microfluidic workflow used in this paper in schematic form (see Supplementary Figure S3, SI, for the detailed chip design). Droplets containing CAZyme-expressing cells are generated in a flow-focussing device. The cells are grown in droplets, express the encoded CAZymes, followed by a picoinjection with a lysis solution that also contains the reagents for subsequent coupling reactions. Finally, the droplets are sorted based on the reaction progress (measured by the emergence of an absorbance signal) within a defined incubation period.

**Figure 3:**
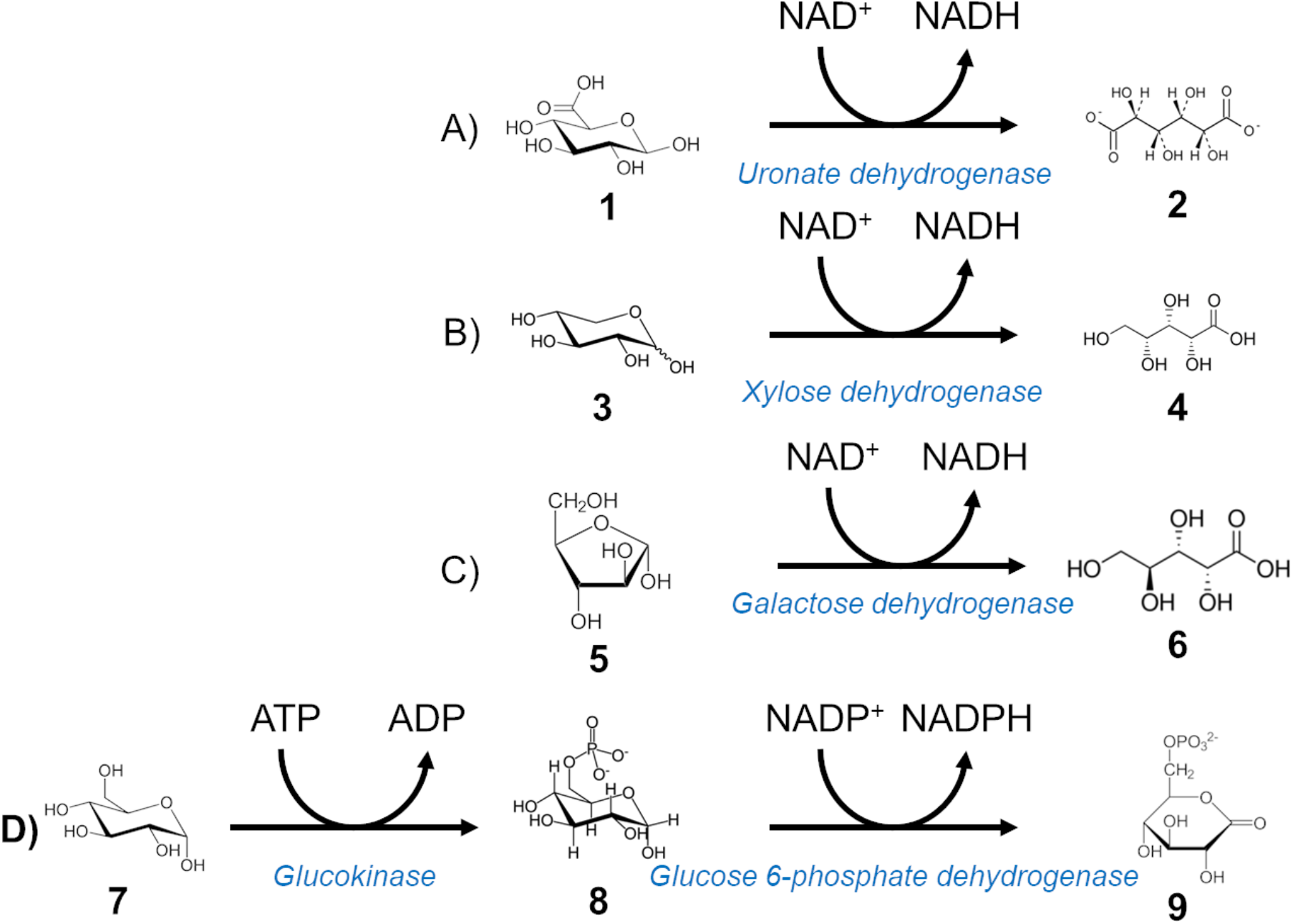
Figure 3. Detection of downstream reaction products. (**A**) Coupling reaction for detection of glucuronic acid. Glucuronic acid (**1**) is converted to glucaric acid (2) by uronate dehydrogenase with a concomitant NADH release. (**B**) Xylose (**3**) conversion to xylonic acid (**4**) by Xylose dehydrogenase. (**C**) Coupling reaction for arabinose detection. Galactose dehydrogenase releases NADH whilst converting L-arabinose (**5**) to L-arabinonic acid (**6**). (**D**) Multi-enzyme cascade of reactions for glucose detection. Glucose (**7**) is first converted to glucose-6-phosphate (**8**) by glucokinase using ATP, then to 6-phosphoglucono-δ-lactone (**9**) by glucose-6-phosphate dehydrogenase with concomitant generation of NADPH.

## RESULTS

### Validation of downstream product detection by a reaction cascade

A robust selection system using coupling reactions (i) must ensure that the coupling reaction is not the rate-limiting step, and (ii) requires highly specific coupling enzymes for the monosaccharide released in the reaction of interest to avoid false positive signal. This is particularly important in the context of a complex natural substrate that contains various types of sugars. Our objective is a versatile workflow in which one could switch from one target activity to another by simply using a different coupling enzyme to detect breakdown of a different natural substrate in an identical technical setup (defined by microfluidic chip designs, scripts & instruments, reagent volumes, on- or off-chip incubation procedures). Commercially available enzymes gave us the opportunity to focus on the detection of xylose, arabinose, glucuronic acid and glucose as examples. The conversion of the first three was achieved in one step, using the enzymes xylose dehydrogenase, uronate dehydrogenase and galactose dehydrogenase, respectively. They convert xylose, glucuronic acid, and arabinose to xylonic, glucaric and arabinonic acids, respectively, together with the concomitant generation of NADH. Glucose by contrast is detected through a 2-step cascade involving the enzymes glucokinase and glucose-6-phosphate dehydrogenase, as it first needs to be activated to glucose-6-phosphate before conversion to 6-phosphoglucono-δ-lactone with concomitant release of NADPH. All enzymatic cascades were tested for substrate specificity by incubating the enzymes with a selection of pure monosaccharides commonly found in (or associated with) the degradation of various plant polysaccharides (glucose, mannose, xylose, arabinose, glucuronic acid, galactose and fucose). The results confirmed that all coupling systems except galactose dehydrogenase were at least an order of magnitude specific for their respective target monosaccharides (Supplementary. Figure S1, SI). The pair glucokinase/G6PDH and uronate dehydrogenase showed no detectable signal when incubated with other monosaccharides (*i*.*e*. different than glucose and glucuronic acid, respectively), suggesting a preference of >20-fold above background. Xylose dehydrogenase showed an activity on glucose equivalent to 5% of that on xylose (after 60 minutes), confirming previously reported values^25^ and indicating a 20-fold specificity. Galactose dehydrogenase was the only enzyme with relevant promiscuity, showing similar activities on galactose and arabinose, in agreement with a previous report^26^. Even so, this enzyme can still be effectively used for the detection of arabinose released by arabinofuranosidases (Ara*f*ases) from beechwood xylan (BX) or wheat arabinoxylan (WAX), as they are devoid of galactose. On the other hand, it would be unsuitable for screening enzymes acting on a substrate containing both galactose and arabinose (*e*.*g*. corn glucuronoarabinoxylan, eucalyptus xylan, arabinogalactan).

To demonstrate the biological relevance of these coupled assays in the context of natural substrate degradation, these cascades were validated in multiwell plate format on natural polysaccharides (cellulose, beechwood xylan and wheat arabinoxylan) using purified CAZymes to release specific monosaccharides from plant heteropolymers.

The coupling enzymes were first assayed on various monosaccharides for their specificity (Supplementary Figure S1) and the results validate the use of these coupling reactions to quantitatively detect cellulase, β-glucosidase, xylanase, β-xylosidase, arabinofuranosidase and glucuronidase activities when assayed on natural substrates. Supplementary Figure S2 shows the synergistic effects observed upon addition of accessory CAZymes (*e*.*g*. β-glucosidase, β-xylosidase, α-glucuronidase, etc.) to the primary CAZymes (*e*.*g*. xylanase, cellulase, etc) for the multistep deconstruction of linear or branched plant polysaccharides.

### Assay miniaturisation to the droplet scale

In order to demonstrate the applicability of our detection system in a 10^6^-fold scale down format in picoliter droplets, we generated two droplet populations using the purified enzymes used in the plate assays in a flow-focussing chip design for parallel droplet production (Supplementary Figure S3A). These two populations contained either the primary targeted CAZyme activity (β-glucuronidase, cellulase, endo-xylanase or ara*f*ase) and the corresponding natural substrate, or served as a negative control population containing only the substrate without the first CAZyme in the cascade. The reaction product was monitored with an absorbance sorter. In contrast to the plate experiments described above, a coupled reaction leading to a more strongly absorbing product (via an electron carrier, 1-methoxy-5-methyl-phenazinium methyl sulfate (mPMS) that transfers electrons from NAD(P)H to the terminal electron acceptor dye (Figure 1; also used for NAD^+^-dependent amino acid dehydrogenases^5^) was used instead of directly monitoring NAD(P)H at 340 nm. The detection reaction generates a reduced ‘Water-Soluble Tetrazolium-1’ (WST-1), a dye absorbing at 455 nm with a molar extinction coefficient higher than NADH (3.7 × 10^4^ at pH8 vs 6.2 × 10^3^ M^−1^.cm^−1^)^27–29^ that makes the reaction 6-fold more sensitive.

Both droplet populations were reinjected on a pico-injection chip (Supplementary Figure S3D) to fuse each droplet to a defined volume of the respective coupling reaction mix (containing coupling enzyme, cofactors, buffer). To allow WST-1 production, these droplets were further incubated for 1h at the optimal temperatures of the respective coupling enzyme(s) in custom-made oil-filled containers^24^ before reinjecting the droplets into a chip for Absorbance-Activated Droplet Sorting (AADS) (Supplementary Figure S3E).

A distribution plot of the absorbance of the combined droplet populations (Figure 4A-D) shows a clear bimodal distribution between those with and without the first CAZYme of the cascade. In the conditions of the assay, we estimated that a 0.005 absorbance value between the control peaks seemed sufficient to accurately sort out the positive droplets. For β-glucuronidase and ara*f*ase activities, when assayed on aldouronic acids and WAX, the signal differences ΔA between positive and negative controls (normalized to a 0-absorbance value) were 0.091 and 0.044, respectively. When assayed alone on BX, the GH10 xylanase generated a 0.075 absorbance difference with the negative control. The combination of this enzyme with a GH43 β-1,4-xylosidase was enhancing the reaction, as the xylose amount effectively released in this case corresponded to a doubling (0.143) absorbance difference with the negative control. To confirm the synergistic effects observed in plate assays using the GH10 xylanase and the GH51 ara*f*ase on WAX and to demonstrate the ability of the system to quantitatively measure the activities of the target enzyme, a triple flow-focussing chip design was developed, allowing to generate three populations of droplets in parallel (Supplementary Figure S1B). This set up was also used to demonstrate the synergistic activity of cellulase and β-glucosidase to release glucose from Carboxymethyl-Cellulose (CMC). The three populations contained either a negative control (no enzyme), a positive control (xylanase alone/cellulase alone), or a synergetic control (xylanase + ara*f*ase / cellulase + β-glucosidase). As shown in Figure 4E&F, an additional droplet population associated with the synergetic condition could be detected, generating in both cases a 100% increase in absorbance signal difference with the negative control. Given that the endoglucanase used here is only able to generate very small amounts of glucose from CMC, it is not surprising to find that β-glucosidase supplementation is required to ensure a clear separation of peaks. Taken together these results demonstrate the capacity of the screening strategy to efficiently and specifically detect the desired CAZyme activities in droplets.

**Figure 4:**
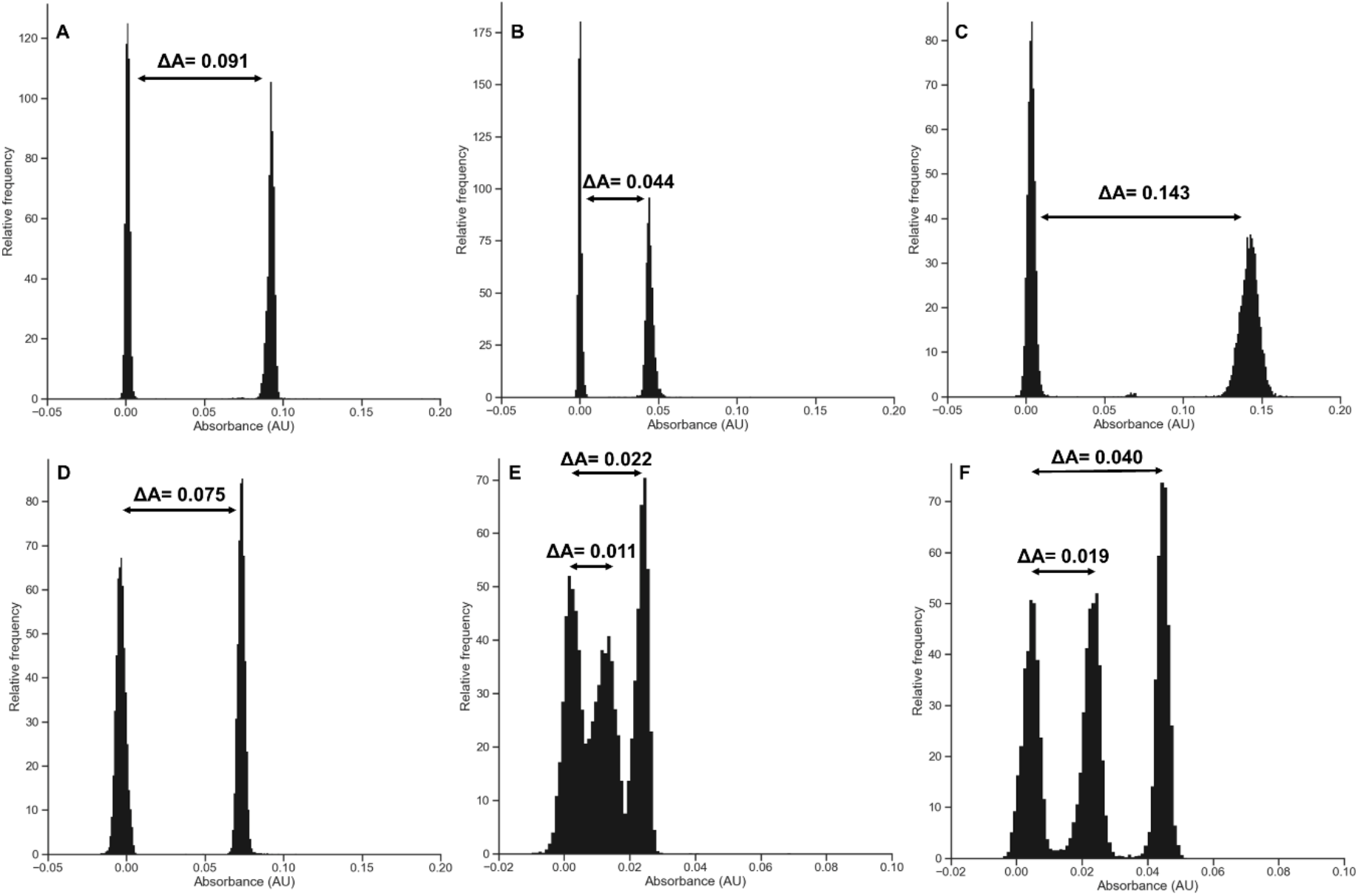
Development of NADH-based coupled assays in droplets for the detection of CAZyme activity on natural substrates. As coupling enzymes were added in excess, signal differences reflect CAZymes’ intrinsic specific activities. Two control populations of droplets of identical size containing either the substrate alone in buffer (AldoUronic acids (**A**), Wheat arabinoxylan (**B**) or Beechwood xylan (**C, D**)) or supplemented with GH67 glucuronidase (**A**), GH51 Arafase (**B**), GH10 xylanase + GH43 β-xylosidase (**C**) or GH10 xylanase alone (**D**) were generated using the double flow-focussing droplet generator shown in Supplementary Figure S3A. Three control droplet populations containing either CMC alone in buffer, supplemented with GH5 endo-cellulase, or supplemented with GH endo-cellulase + GH1 β-glucosidase (**E**), wheat arabinoxylan alone in buffer, supplemented with GH10 xylanase alone, or with GH10 xylanase + GH43 β-xylosidase (**F**) were generated using the droplet generator shown in Supplmentary Figure S3B. The droplets were further picoinjected with uronate dehydrogenase (**A**), galactose dehydrogenase (**B**), xylose dehydrogenase (**C, D, F**) or glucokinase + G6PDH (**E**) together with the appropriate cofactors, mPMS and WST-1 using the picoinjector shown in Supplementary Figure S3D. After a 1h incubation at 37°C (**A, B, C, D, F**) or 25°C (**E**), the droplet absorbance was measured using the sorter shown in Supplementary Figure S3E.

### Demonstration of activity in droplets using cellular lysates

The next step in the development of our screening platform was to demonstrate the ability to detect positive hits when activity levels correspond to actual expression levels of CAZymes found in cellular lysates. Five suitable candidates for *E. coli* expression allowing us to assess the various CAZyme activities with purified enzymes were identified in the literature: we used the *Clostridium thermocellum* GH5E, a bi-functional cellulase/xylanase^30^, the Araf62A ara*f*ase from *Streptomyces thermoviolaceus* OPC-520^31^, the *Humicola insolens* GH43A β-xylosidase^32^, the *Paenibacillus barengoltzii* Xyn10A xylanase^33^ and the GH67 glucuronidase from *Dictyoglomus turgidum* DSM 6724^34^. In order to allow a comparison of expression levels between these enzymes, all were cloned in the same pET26b vector (to ensure homogeneous expression) and expressed in *E. coli* BL21 (DE3) to assay their activity in droplets. As reported in the literature ^31–33^ all proteins except GH5 members were secreted to some extent. This had consequences on the workflow used for the encapsulation of cells: to achieve the desired droplet occupancy (λ=0.2) when generating the droplets, induced cells had to be diluted in the flow-focussing chip. For the cellulase, the cells were grown in LB medium and induced with IPTG prior to co-encapsulation with the lysis solution, the substrate (CMC) and a β-glucosidase used to convert the cellobiose molecules generated by the GH5 to glucose. After incubation with the substrate, the 100 pL droplets were reinjected on a picoinjection chip to add the other reactions cascade components. After a final incubation to allow WST reduction, the 200pL droplets were reinjected on an absorbance chip for AADS analysis. As shown in Figure 5A, a distinct peak corresponding to cellulase containing droplets could be observed, with a ΔA corresponding to 192 µM WST-1. The four other CAZymes are secreted, so growth and induction prior to droplet encapsulation was impossible, as activity from secreted protein would be present in every droplet. The workflow was therefore slightly modified. Using the same flow focussing device, 100 pL droplets containing non-induced single cells were incubated for 48 h to allow for cellular growth and induction of protein expression in the droplets by co-encapsulation in auto-inducible medium. Then, the cascade components were picoinjected resulting in 200 pL droplets. No cell lysis was required, as the proteins are secreted. However, as WST-1 is a reporter of cell viability, background form cell growth was reduced by addition of the non-lytic spectromycin antibiotic. As shown in Figure 5B-F, even if the signal difference was smaller than with purified enzymes, all cell-containing populations could be differentiated from empty droplets with a ΔA corresponding to 424, 197, 183, 660 and 80 µM WST-1 for *Hi*Xyl43A on BX (B), *Hi*Xyl43A on X2 (C), *Sth*Araf62A on WAX (D), *Pb*Xyn10A on BX and *Dt*GH67 on uronic acids (F), respectively.

**Figure 5:**
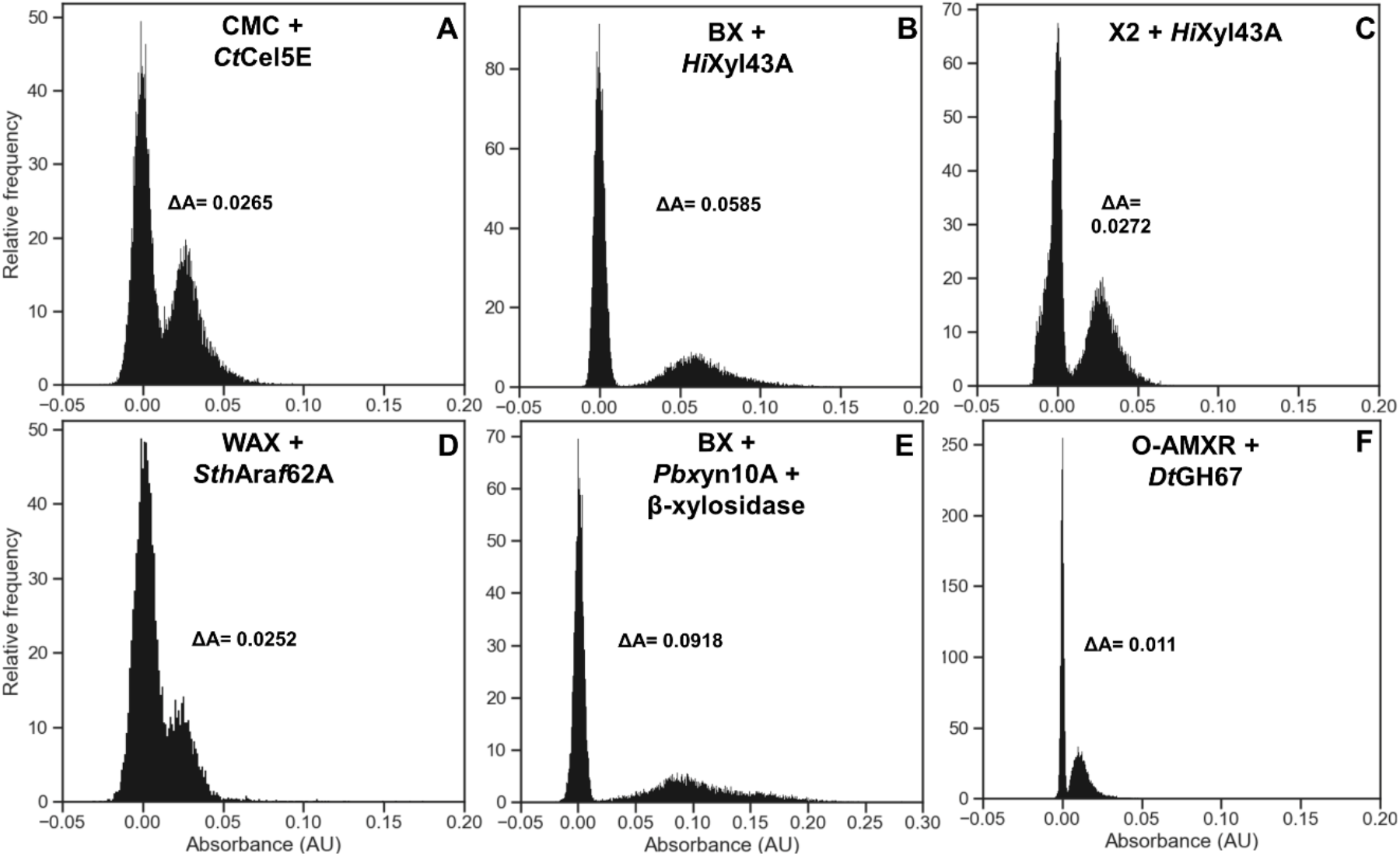
Detection of CAZyme expressing cells in droplets using the coupled assays. Droplets containing *E. coli* BL21 (DE3) cells transformed with plasmids encoding *Ct*Cel5E (**A**), *Hi*Xyl43A (**B, C**), *Sth*Araf62A (**D**), *Pb*Xyn10A(**E**) or *Dt*GH67 genes (**F**) were generated using the flow-focussing device shown in Supplementary Figure S1C. After cell growth and protein expression, these droplets were picoinjected with CMC + glucose cascade, beechwood xylan + xylose cascade, xylobiose + xylose cascade, wheat arabinoxylan + arabinose cascade, beechwood xylan + β-xylosidase + xylose cascade or uronic acids + glucuronic acid cascade, respectively. Droplets were incubated at 25°C for 48h (**A**), 37°C for 2h (**B, C, D, F**) or 37°C for 24h (**E**), prior to absorbance measurement with the sorter shown in Supplementary Figure S1E.

### Functional screening of CAZyme activity from multiple activities containing cell mixtures

The final development step of our CAZyme sorter was to demonstrate the efficiency of the screening platform in achieving accurate selection of the desired activity in a biological context. We therefore designed an enrichment experiment mimicking a biological situation in which the expected hit is rare among a library of cells expressing various other enzymatic activities (Figure 6), and notably other CAZymes. We used β-xylosidase as the activity of interest to establish the proof-of-concept. As 3 out of the 4 used proteins in this assay are secreted, mixing the cells prior to growth and protein expression was impossible. To avoid any bias related to protein secretion, cells expressing *Hi*Xyl43A, *Ct*Cel5E, *Dt*GH67 and *Sth*Araf62A were mixed (1:33:33:33) and directly encapsulated into 50 pL droplets containing auto-inducible medium and cultivated for 48h to induce cell growth and protein expression without risking any cross-contamination. This situation seemed reasonably representative of real-life sorting experiments, as many endo-acting CAZymes are commonly secreted. These ratios would also be representative of a typical library sorting experiment, for which the top 1% of absorbent cells are sorted. To specifically enrich the β-xylosidase expressing droplets, the picoinjection solution was containing xylobiose as natural substrate and the xylose dehydrogenase cascade components to detect the activity. Three types of droplets would be picoinjected at this stage: i) empty droplets, ii) droplets containing *Ct*Cel5E expressing cells (not secreted) and iii) droplets containing one of the three other CAZymes expressing cells (secreted). Here again, to avoid any possible bias in the selection related to protein secretion, we decided to uniformly lyse all the cells and therefore added a lysis agent in the picoinjected solution. The WST-1 molecule used for detection is a dye very well known for being reduced by a multitude of cellular oxidoreductases^35^. To reduce background absorbance, we decreased the ratio of cell lysate to picoinjected solution down to 1:3 by picoinjecting 150 µL of reagents into 50 pL droplets, ensuring a greater detection specificity. The picoinjected solution also contained pyranine as absorbance offset and 1-Bromo-3,5-bis(trifluoromethyl)benzene as refractive index modifier to ease the peaks detection by the in-house Arduino script used for droplet detection and absorbance measurement. Nine hundred twenty-eight 200 pL droplets were sorted from 1.28 million (72% of expected positive droplets number) at an average frequency of 175 Hz. The DNA they contained was extracted and transformed into *E. coli* BL21 cells, further plated onto kanamycin-containing agar plates. Ninety-six randomly selected colonies were selected for cultivation into auto-inducible medium, allowing the expression of genes harboured by any kanamycin-resistant plasmid they would contain. As *Hi*Xyl43A is a secreted enzyme, we only assayed the supernatants from the 48h cultures in a secondary screening to confirm the presence of xylosidase activity. Using the same assay conditions as in the plate assay controls, 83 out of the 96 cultured clones (86.5%) generated a high absorbance signal when assayed, both similar in amplitude and kinetics as the positive control, confirming *Hi*Xyl43A presence (Supplementary Figure S4), resulting in an enrichment of 86-fold (calculated according to Zinchenko *et al*.^36^) or 638-fold (Baret *et al*.^37^, Table 1).

**Table 1.** Summary of observed enrichments in droplet assays.

| $\lambda$ | $\epsilon_0$ | Number of droplets | | Number of clones | | Baret *et al.*<sup>37</sup> | | Zinchenko *et al.*<sup>36</sup> | |
| --- | --- | --- | --- | --- | --- | --- | --- | --- | --- |
| | | Analysed | Sorted | *HiXyl43A* | Negative | $\epsilon_1$ | $\eta$ | $\epsilon_1'$ | $\eta'$ |
| 0.1 | 0.01 | 1280000 | 928 | 83 | 13 | 6.38 | 638 | 0.865 | 86.5 |

**Figure 6:**
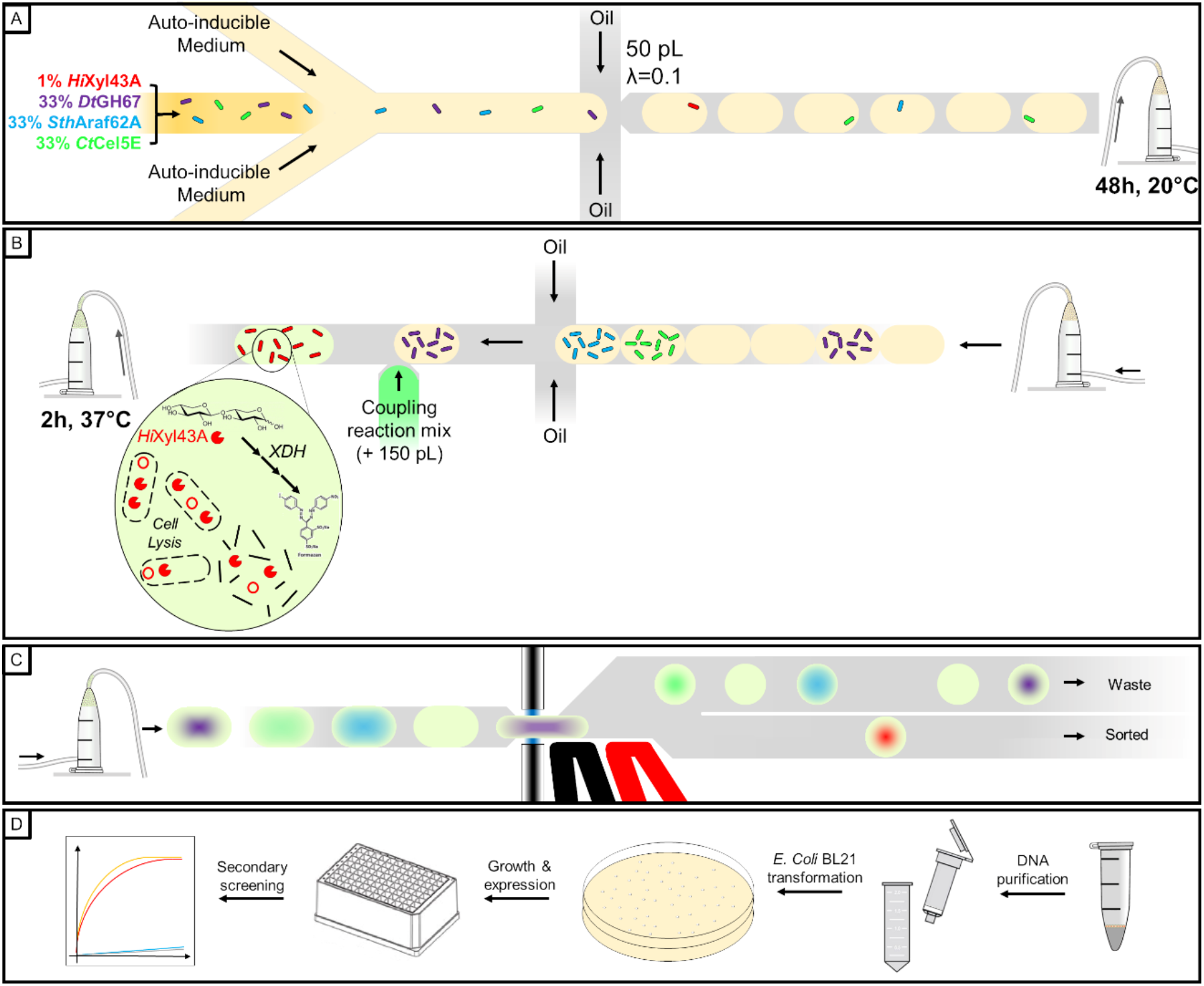
Functional screening workflow of xylosidase activity in mixtures of droplets expressing two different CAZymes. (**A**) Cells expressing *Hi*Xyl43A, *Dt*GH67, *Sth*Araf62A and *Ct*Cel5E were mixed to a 1:33:33:33 ratio and encapsulated into 50 pL droplets with λ=0.1. (**B**) After a 48h incubation allowing cell growth and protein expression the droplets are picoinjected with a 150 pL coupling reaction mix allowing cell lysis and detection of xylose activity. (**C**) After a 2h incubation allowing signal development, the droplets are sorted by AADS. **D**) The positive droplets are collected, the DNA they contain extracted and purified, cloned into *E. coli* BL21 cells and grown on selective medium. 96 individual colonies are grown in auto-inducible medium allowing protein expression. After centrifugation of the induced cells, the supernatant is assayed for xylose release from xylobiose in a microtiter plate.

### Conclusion and future perspectives

Droplet microfluidics has the potential to make ultrahigh-throughput screening much more accessible and affordable, by replacing the classical test tube with an emulsion compartment and manual or robotic handling with straightforward devices and workflows^38^. Nevertheless, directed protein evolution or metagenomic screening in this format is only possible, when an assay is available that leads to an optical readout. The quality of the selection output is determined by the extent to which the assay substrate resembles the desired function (‘*you get what you screen for’*)^39^. Current ultrahigh-throughput screening of CAZYmes relies on substrates with direct attachment of a fluorophore to a sugar and reaction progress is measured by its hydrolytic release. While these substrates are related to natural sugars, they are not identical in structure and reactivity and might lead screening efforts astray. In addition, such substrates are generated by means of organic synthesis that has to be redeveloped for every sugar substrate construct, making it difficult for researchers without these skill sets who need to rely on commercial availability.

Here, we demonstrate that the degradation of natural substrates can be monitored indirectly through a coupled reaction: a specific primary assay can be translated by a subsequent reaction into an optical signal in a unified way for multiple substrates, so that the primary enzyme can be modularly exchanged and still be detected by the same secondary assay.

We demonstrated that it is possible to detect the release of four types of the most common monosaccharides found in natural hetero-oligosaccharides or -polysaccharides produced by specific CAZyme activities in droplets. We then showed that these coupling reactions are effective under biological conditions, with model enzymes expressed from plasmids, and both secreted or intracellular enzymes are amenable. The enrichment experiment of a β-xylosidase from a mixture of cells expressing three other different CAZyme activities, with an 86.5% proportion (from an initial abundance of 1% in the cell mixture) illustrated the specificity of this system. This coupled reaction strategy now expands the potential of droplets microfluidics to large-scale functional screening of CAZymes.

Previous efforts to use cascades of reactions to detect CAZymes in droplets remained limited^11^: cellulase activity was monitored by hydrogen peroxide production using hexose oxidase, a non-selective enzyme (with activity against various monosaccharides and oligosaccharides^40^). This promiscuity of the coupling enzyme would lead to a non-specific response when assaying natural polysaccharides containing various monomer types, as it is frequently the case in biomass degradation. The coupling reaction generates hydrogen peroxide that is then monitored by production of fluorescein from the expensive probe AminoPhenyl Fluorescein (APF). Vanadium bromoperoxidase, the enzyme required for this conversion, is an allosterically regulated enzyme with narrow H_2_O_2_ dynamic range due to bleaching, which might become the rate-limiting step in the cascade of reactions. The final claimed enrichment factor of 300, actually corresponds to 30% positive clones after sorting from a 0.1% cellulase-containing initial library, meaning that 70% false-positives are retrieved from the sorting of an activity present at typical abundance in a library.

Similarly, the detection of cellulase activity^41^, or β-glucosidase, cellobiohydrolase-I, and endoglucanase activities^42^ were also based on the unspecific reaction of hexose oxidase and made use of ionic liquid solubilized native cellulose (Avicel) or biomass (Switchgrass) as substrates. In both cases, the elemental monosaccharide detected is glucose, limiting the range of screenable CAZy activities that can be addressed.

In order to expand the catalogue of addressable reactions, new specific coupling reactions had to be identified. The use of monosaccharide-specific dehydrogenases ensures a substantial specificity for the actual natural product of the reaction, even in the case of a natural substrate formed of various types of carbohydrates. Several of these dehydrogenases are commercially available and produce a similar by-product (NADH or NADPH), so that detection is uniform and would not need to be tailored for each reaction. On the other hand, specificity for monosaccharide means that only *exo*-CAZymes that remove one glycoside unit at a time can be assayed. For the detection of *endo*-CAZyme activity, coupling enzymes that accept the respective fragment produced or addition of enzymes to break down fragments further into monosaccharides would be necessary.

The modular architecture of the microfluidic set-up makes it possible to adjust the workflow: as demonstrated here by the addition of coupling reagents (deliberately at a stage when the reaction is considered complete) and a period of cell growth; or in the future by allowing brief on-chip vs. long-term off-chip incubation of droplets, changing reaction conditions (by adding reagents or changing pH) or timed addition of lysate reagents. The modularity will generally help to overcome chemical incompatibilities of target and coupling enzymes. More complex multistep cascades with different chemical transformations can be envisaged when single step cascades are impossible. For example, aforementioned coupling to sugar oxidases^11,43,44^ could be carried out under conditions where the detection enzyme does not become limiting. Carbohydrates functionalised *e*.*g* with sulfate, amine or acetate groups could be detected through cascades that monitor their release (*e*.*g*. of phosphate by sugar phosphorylases monitored with the molybdenum blue assay^45^) or by direct detection of phosphorylated sugars^28^. Moving beyond catabolic CAZymes, glycoside transferases that consume NADH may become detectable with a lactate/pyruvate dehydrogenase recycling system (again monitored by the release of H_2_O_2_)^46^.

While a large number of potential cascades for CAZyme reactions remain to be established (beyond the four types of monosaccharides shown here), the modular workflow will enable implementation of various strategies that will expand the scope of ultrahigh throughput screening in droplets to large-scale discovery campaigns of useful and high performing CAZymes.

## Materials and Methods

### Chemicals, substrates, enzymes and plasmids

Monosaccharide standards (xylose, glucuronic acid, arabinose, glucose, galactose and fucose), CarboxyMethyl-Cellulose, Avicel PH-101, NAD^+^, NADP^+^, ATP, MgCl_2_, cellobiose, 1-Methoxy-5-methylphenazinium methyl sulfate (mPMS, ref M8640) were purchased from Sigma Aldrich. Xylobiose (O-XBI), Cellohexaose (O-CHE), Aldouronic acids mixture (O-AMXR), Beechwood xylan (P-XYLNBE), Wheat arabinoxylan (P-WAXYL), endo-β-1,4-glucanase (E-CELBA), β-glucosidase (E-BGLUC), a mixture of galactose dehydrogenase and galactose mutarotase (E-GALMUT), uronate dehydrogenase (K-URONIC), α-glucuronidase (E-AGUBS), xylose dehydrogenase (K-XYLOSE), β-xylosidase (E-BXEBP) and endo-β-1,4-xylanase (E-XYLNP) were purchased from Megazyme, Ireland. Water Soluble Tetrazolium salt (WST-1, ref W201-10) was purchased from NBS Biologicals, Huntingdon, UK, while rlysozyme was sourced from Merck and 1-Bromo-3,5-bis(trifluoromethyl)benzene from Alfa Aesar. Glucokinase (AE00171), Glucose-6-phosphate dehydrogenase (AE00111), Arabinofuranosidase 51B from *Cellvibrio japonicus* (CZ0707) were purchased from NZYtech, Lisboa, Portugal.

### Polysaccharide preparation

A hundred milligrams of beechwood xylan and wheat arabinoxylan were solubilised in 0.5 mL of 95% EtOH before addition of 9.5ml of ddH2O. The slurries were heated on a hot plate for 4h at 95°C under magnetic stirring and left O/N in a 65 °C oven. After dry weight measurements, final concentrations of 0.85% and 0.47% (w/v) were achieved for beechwood xylan and WAX, respectively. A 1% CarboxyMethyl-Cellulose stock solution was prepared by solubilising 100 mg of CMC into 10 mL of hot water and incubating the solution for 24 hours at 65°C. These substrates were aliquoted and stored at −20°C.

### Determination of the specificity of the coupling enzymes

One millimolar of xylose, glucuronic acid, arabinose, glucose, galactose, mannose and fucose standards were incubated with each enzyme or mixture of enzymes (Uronate dehydrogenase, xylose dehydrogenase, galactose dehydrogenase / galactose mutarotase) in 100mM sodium phosphate pH 7.0 for 1h at 25°C in presence of 1 mM NAD^+^. Enzymes were added according to manufacturer’s recommendations and assayed in triplicates (2 µL of enzyme solution for 200 µL reaction volume). For assessing the specificity of the glucose coupled assay, 1 mM of the same monosaccharides were incubated with 0.1 U of glucokinase and 1 U glucose-6-phosphate dehydrogenase, supplemented with 10 mM ATP, 1 mM MgCl_2_ and 1 mM NADP^+^ in 100 mM sodium phosphate pH 7.0 for 1h at 25°C. A negative control was added for normalisation, consisting of each monosaccharide, buffer, ATP, MgCl_2_ and NAD(P)^+^ in concentrations equivalent to the assays with enzymes. Absorbance at 340 nm was recorded on an infinite M200 plate reader (Tecan) using µCLEAR® black 96-well plates (Ref 655906 Greiner). These assays were performed in triplicates.

### Controls for the detection of CAZyme activities in coupled reactions

A final concentration of 0.1% (w/v) of beechwood xylan, wheat arabinoxylan or 1 mM xylobiose was incubated in 100 mM sodium phosphate pH 7.0 in presence or absence of 10 units of GH11 xylanase (E-XYLNP), 0.11 units of GH43 β-1,4-xylosidase (E-BXSEBP), 0.09 units of GH67 α-glucuronidase (E-AGUBS) and 0.25 µg of *Cj*Abf51B Arafase (CZ0707). These enzymes mixes were supplemented with 1 mM of NAD^+^ and xylose dehydrogenase (2 µL of commercial solution, as recommended by the manufacturer). Similarly, 0.1% (w/v) of beechwood xylan, wheat arabinoxylan and aldouronic acids mixture (O-AMXR) were incubated in 100 mM sodium phosphate pH 7.0 with or without 0.09 units of GH67 α-glucuronidase (E-AGUBS), 10 units of GH11 xylanase (E-XYLNP) and 0.11 units of GH43 β-1,4-xylosidase (E-BXSEBP). To allow quantification of glucuronic acid, these enzyme mixes were supplemented with 2 µL of uronate dehydrogenase and 1 mM NAD^+^. Beechwood xylan or wheat arabinoxylan were assayed for arabinose release. Ten units of GH11 xylanase (E-XYLNP) with or without 0.11 units of GH43 β-1,4-xylosidase (E-BXSEBP), or 0.25 µg of *Cj*Abf51B arabinofuranosidase (CZ0707) were incubated with these substrates at a final concentration of 0.1% (w/v). As only the β-form of l-arabinose and d-galactose is recognised by β-galactose dehydrogenase and because of the low rate of natural mutarotation between the α- and β-anomers, the commercial solution of galactose dehydrogenase is supplemented with galactose mutarotase to get a rapid mutarotation to galacturonic and arabinonic acids. Two microlitres of this commercial solution of coupling enzymes were added together with 1 mM of NAD^+^ to the enzyme mixes. Glucose was detected from cellulose degradation. A final concentration of 0.1% (w/v) Avicel PH-101, CMC or 1 mM cellohexaose was incubated with 1.2 units of GH5 cellulase (E-CELBA) with or without 0.04 units of GH3 β-1,4-glucosidase (E-BGLUC) in 100 mM sodium phosphate pH 7.0. These mixes were supplemented with 0.1 units of glucokinase and 1 unit of glucose-6-phosphate dehydrogenase in presence of 1 mM NADP^+^, 10 mM ATP and 1 mM MgCl_2_. These coupling reaction assays were incubated for 1h at 25°C in a final volume of 200 µL and made in triplicates. Absorbance at 340nm was recorded on an infinite M200 plate reader (Tecan) using µCLEAR^®^ black 96-well plates (Ref 655906 Greiner).

### Chip design and microfluidic device fabrication

Chip designs can be downloaded from our open access repository DropBase (https://openwetware.org/wiki/DropBase:Devices as .dwg & .png files). The designs for the poly(dimethyl)siloxane (PDMS) chip devices were prepared using AutoCAD software and designs are shown in Supplementary Figure S3. The devices were fabricated on 3-inch silicon wafers (MicroChemicals) with standard soft lithographic procedures by using a high-resolution acetate mask (Microlithography Services Ltd.). The photoresist material SU-8 2050 and SU-8 2075 was used to obtain a 50 µm channel height and 80 µm for multi-flow focussing chips and sorter, respectively (Supplementary Figure S3A, B & E). The resulting lithographic master moulds were used to generate polydimethylsiloxane (PDMS) chips by pouring 20 g of PDMS monomer and curing agent at a ratio 10:1 in a Petri dish (Ø: 9 cm). After degassing and PDMS solidification (65°C, 4h), PDMS was activated by exposure to an oxygen plasma system (Femto, Diener electronic) and devices were bonded to a microscope glass slide. Hydrophobic modification of the channels surface was achieved by injecting a solution of 1% (v/v) trichloro (1H,1H,2H,2H-perfluorooctyl) silane in HFE-7500 oil (3M Novec) into the channels. Methods for fabrication of absorbance sorting chips are detailed in Gielen *et al*^5^.

### Sorting electronics

The voltage signal from the photodetector was split in two, recorded with a custom LabVIEW program [using an Analog-to-Digital Converter (National Instruments, USB-6009)], and at the same time sampled in 12-bit resolution via an analog-in pin of a 32-bit Cortex M3 ARM-based microcontroller (Arduino Due). To match the voltage of the detector (max., 10 V) to the maximum voltage tolerable for the microcontroller board (3.3 V), a simple voltage divider (10 kΩ resistors) was used. On the microcontroller, the signal from the detector was compared with a threshold value for sorting. Below the threshold, a pin was set high to activate a pulse generator (TGP110, Aim-TTi), used to generate a 6.5-V pulse, and manually adjusted to 10 kHz, in order to be smaller than the droplet period (5 ms at 200 Hz). The pulse generator controlled a function generator (20 MHz DDS Function Generator TG 2000, Aim-TTi) working in external gated mode, generating a 10-kHz square signal at an adjustable amplitude, which was then amplified 100 times with a voltage amplifier (TREK 601c) to actuate the sorter. This setup enabled convenient manual adjustment of all important sorting parameters such as droplet size and voltage gates (see SI).

### Controls for coupling reactions in droplets

Two or three populations of identical size droplets were generated using the co-flow focussing devices shown in Supplementary Figure S3A&B by running the aqueous phases with HFE7500 supplemented with 2.5% (w/v) 008-FluoroSurfactant (RAN Biotechnologies) at flow rates of 4 µL.min^−1^ and 20 µL.min^−1^, respectively. The chips were operated with syringe pumps (Nemesys, Cetoni) and 100 µL (for aqueous phases) or 1 mL (for oil) gas-tight glass syringes (SGE). Positive populations contained the substrate (1 mM of xylobiose or 0.1% w/v of wheat arabinoxyan, beechwood xylan, CMC or aldouronic acids) and CAZymes (various combinations of xylanase, β-1,4-xylosidase, arabinofuranosidase, α-glucuronidase, cellulase and β-1,4-glucosidase in concentrations identical as for the plate controls) in 100 mM sodium phosphate pH 7.0, supplemented with 1 mg.ml^−1^ BSA. Negative droplet populations contained the same mixture but lacked the CAZymes. The droplets generated in this way were collected in custom-made collection containers consisting^47^ of a 0.5 mL Eppendorf tube glued upside down on a microscope glass slide through a ∅ 0.8mm (I.D.) PTFE tubing (Bola Bohlender) or ∅ 0.86mm PE (Smith Medical) connected from the top and filled with oil. An additional tubing was connected at the bottom of the tube in order to avoid over-pressure during the collection, and to allow re-injection of the droplets during the subsequent steps. Upon generation, these droplet populations mixtures were reinjected on a pioinjection chip (Supplementary Figure S3D) at 4 µL.min^−1^ with a 4 µL.min^−1^ spacing oil flow. Appropriate coupling reaction reagents containing identical concentrations of coupling enzymes as for the plate assays, supplemented with 1 mM NAD^+^, 5 µg.mL^−1^ mPMS and 1 mM WST-1 were picoinjected at a flow rate of 1 µL.min^−1^. For glucose detection, NAD^+^ was replaced by NADP^+^, and 10 mM ATP plus 1 mM MgCl_2_ were added to the mix. Droplet coalescence was induced by generating 2.5 V, 10 kHz square waveform using a 20 MHz DDS function generator TG 2000 (AimTTi). The signal was then amplified 100 times using a high voltage amplifier (TREK 601c) and applied onto the chip through 5 M NaCl-filled tubing connected to the electrode channels. A 1h-room temperature incubation time was applied to allow WST-1 conversion before analysis on the absorbance chip shown in Supplementary Figure S3E. The droplets were reinjected onto the sorting chip at a flow rate of 2 µl.min^−1^ with a flow of spacing oil without surfactant of 20 µl.min^−1^. As the droplets were not collected in analysis mode, a similar setup to the one used for sorting was used, but with the Arduino board turned off. Data was recorded with a custom LabVIEW program, and processed with home-made Python scripts. Overall, the total incubation time of CAZymes with the substrate from aqueous phase preparation to measurement was 4h at room-temperature.

### Cell controls in droplets

*E. coli* codon-optimized genes encoding for a GH5 (cellulase catalytic domain from *Hungateiclostridium thermocellum* ATCC 27405, *Ct*Cel5E, residues 290-654)^30^, GH67 *Dictyoglomus turgidum* DSM 6724 α-glucuronidase^34^ (*Dt*GH67), GH62 *Streptomyces thermoviolaceus* OPC-520 α-l-arabinofuranosidase (*Sth*Araf62A)^31^, GH43A *Humicola insolens* Y1 β-xylosidase / α-l-arabinofuranosidase (*Hi*Xyl43A)^32^ and GH10 Xyn10A from *Paenibacillus barengoltzii* CAU904 xylanase (*Pb*Xyn10A)^33^ were ordered from GeneArt (ThermoFisher) and cloned into pET26b expression vector with a C-terminal hexahistidine tag by the Gibson assembly method. All sequences except *Ct*Cel5E contained the native signal peptide. *E. coli* BL21 (DE3) cells harbouring these various plasmids were individually encapsulated in autoinducible medium (AIM, [NZYtech, Lisboa, Portugal]) supplemented with 50 µg.mL^−1^ kanamycin using the flow focussing device shown in Supplementary Figure S3C. The aqueous phase was composed of a binary mixture (5 µL.min^−1^ each) of cells and fresh autoinducible medium supplemented with 50 µg.mL^−1^ kanamycin was and injected at a flow rate of 10 µL.min^−1^ from two separated inlets. 100 pL droplets were generated at ca. 1.6 kHz from this 2-fold dilution by running the aqueous phase along an 18 µL.min^−1^ HFE7500 supplemented with 2.5% RAN surfactant oil phase, resulting in single cell encapsulation at λ= 0.2 from a starting cell OD of 0.008. The droplets were collected in custom-made containers and subsequently incubated for 48h at 20 °C to allow cell growth and protein expression. Droplets were re-injected on the picoinjection chip (Supplementary Figure S3D) at ca. 500 Hz using a flow rate of 3 µL.min^−1^ with a 5 µL.min^−1^ flow of spacing oil containing 2.5% surfactant. Appropriate coupling reaction reagents were picoinjected at a flow rate of 3 µL.min^−1^ to the droplets, resulting in addition of 100 pL of solution to each droplet. The coupling mixes composition were identical to the droplet controls with pure enzymes. As *Hi*Xyl43A, *Dt*GH67 and *Sth*Araf62A are secreted enzymes, the mixes were supplemented with 100 µg.mL^−1^ streptomycin to reduce noise resulting from cell growth, while 0.1 mg.mL^−1^ polymyxin B and 30 U/mL rLysozyme were added for lysis of the intracellular *Ct*CelE5.

### Microfluidic enrichment experiments

Four *E. coli* BL21 (DE3) precultures, freshly transformed with plasmids encoding *Hi*Xyl43A, *Ct*Cel5E, *Dt*GH67 and *Sth*Araf62A were separately grown in LB medium prior to mixing. After an O.D.600nm > 3.0 was measured, *Hi*xyl43A expressing cells were diluted at a ratio of 1:100 among *Ct*Cel5E, *Dt*GH67 and *Sth*Araf62A expressing cells in autoinducible medium (NZYtech) supplemented with 50 µg.ml^−1^ kanamycin, and the cell concentration was adjusted to an O.D.600nm of ~0.008. This mixture of cells was injected into the flow focussing device shown in Supplementary Figure S3C at a flow rate of 5 µL min^−1^ using a 1 mL SGE syringe. At the chip junction, this channel supplying the cell suspension was mixed with another aqueous solution only containing fresh autoinducible medium and kanamycin at a flowrate of 5 µL min^−1^. Droplets with a volume of ~50 pL were generated at 1.35 kHz from this 2-fold dilution by running a 24 µL min^−1^ HFE7500 oil phase (supplemented with 2.5% RAN surfactant), resulting in single cell encapsulation at λ=0.1. Droplets were subsequently collected and incubated in a home-made collection chamber for 48 hours at 20°C to allow cell growth and protein expression. Then, droplets were reinjected on the picoinjection chip (Supplementary Figure S3D) at a flow rate of 1.5 µL min^−1^. Spacing oil (containing 2.5% RAN surfactant) was injected at a flow rate of 5 µL min^−1^. The coupling solution was picoinjected into the droplets using a flow of 3.5 µL.min^−1^, resulting in the addition of 150 pL of coupling solution to each droplet at a frequency of 340 Hz. The xylose coupling reaction mix was picoinjected, with components concentrations adjusted to take into account the dilution factor resulting from the unequal volumes of cell containing droplets to picoinjected solution (50 pL + 150 pL). After picoinjection, the droplets contained 10 mM NAD^+^, 5 mM WST-1, 5 µg/mL mPMS, 10 mM xylobiose, 0.1 mg.mL^−1^ polymyxin B, 30 U.mL^−1^ rLysozyme, 5 mM pyranine and 1 µL xylose dehydrogenase/ xylose mutarotase (as supplied from megazyme) in sodium phosphate buffer (200 mM, pH 7). To induce droplet coalescence, a 250 V, a 10 kHz electric field was applied. The resulting 200 pL droplets were subsequently incubated for 2 hours at 37°C to allow completion of the coupled reactions. Droplets were reinjected into the sorting chip (Supplementary Figure S3E) at 150–200 Hz (2-2.5 µl.min^−1^ flow rate for the droplet emulsion with spacing oil injected at 30–35 µl min^−1^. The spacing oil was composed of HFE-7500 with 22.5% 1-Bromo-3,5-bis(trifluoromethyl)benzene (Alfa Aesar) that was added to reduce the side scattering of droplet edges^48^. Approximatively 1.28 million droplets were screened, and the 928 most absorbing droplets were sorted (based on the calculation that there was one positive droplet per thousand screened droplets according to a Poisson distribution) and collected in a 1.5 mL LoBind Eppendorf tube.

### DNA recovery and hit validation

A volume of 1H,1H,2H,2H-perfluorooctanol equivalent to the volume of oil collected in the Eppendorf tube was added to break the emulsion. Plasmid DNA was recovered by extracting the mixture three times with 100 µL of nuclease free water (supplemented with 2 µg.mL^−1^ salmon sperm DNA), vortexing for 1 min, centrifuging for 1 min at 14,000 x G on a tabletop centrifuge and combining the aqueous extracts. The aqueous phase was further purified using a DNA clean & concentrator kit (Zymo Research) and eluted in 12 µL elution buffer. Two µL of purified DNA were used for transforming *E. coli* cells (electro-competent *E. cloni* EXPRESS BL21 (DE3), Lucigen) and subsequently plated on LB-agar plates containing 50 µg mL^−1^ kanamycin. Ninety-six randomly selected clones from this transformation were grown in 500 µL AIM supplemented with 50 µg mL^−1^ kanamycin for 48 hours at 20 °C in 96-deep well plates sealed with Breathe-Easier membranes (Sigma). The culture medium was then separated from the cells by centrifugation for 15 minutes at 4500 x G and 10 µL of culture supernatant was assayed for xylobiose lysis using the protocol detailed in Controls for the detection of CAZyme activities in coupled reactions.

## ASSOCIATED CONTENT

### Supporting Information

Considerations regarding the choice of coupling enzymes and assay robustness, practical considerations regarding the sorting operations, coupling enzymes specificities and control reactions, microfluidic chip designs and secondary screening results of sorted clones, microfluidic system setup and practical signal improvement for automatic droplet detection (DOC). This material is available free of charge via the Internet at http://pubs.acs.org.

## Author Contributions

S.L., S.E. and F. H. conceived the study. S.L performed the experimental work with help of T.S.K (microfluidic device design) and P.J.Z (coding of Arduino scripts), drafted the manuscript and prepared the figures. F.H and

S.L wrote the manuscript with input from all authors.

## ACKNOWLEDGMENTS

We thank members of the Hollfelder group for helpful comments and Dr Liisa van Vliet for help with figure editing.

